# P176S polymorphism in Maspin rewires electrostatic interaction that alters Maspin functionality

**DOI:** 10.1101/2022.05.31.494110

**Authors:** Muhammad Ayaz Anwar, Muhammad Haseeb, Sangdun Choi, Kim Kwong-Pyo

## Abstract

Maspin has been known to regress tumors by inhibiting angiogenesis, however, its roles have been reported to be context and sequence-dependent. There are various proteins and cofactors that bind with maspin possibly explaining the conflicting roles of maspin. Moreover, maspin polymorphic forms have also been linked to tumor regression or survival, for instance, maspin with Ser at 176 (maspin-S176) promotes tumor while maspin with Pro at 176 (maspin-Pro176) has opposing roles in cancer pathogenesis. With the help of long molecular dynamic simulation, a possible link between polymorphic forms and tumor progression has been established. First, the maspin is dynamically stable with either amino acid at 176 position, secondly, differential contacts have been observed among various regions, thirdly, these contacts have significantly altered the electrostatic energetics of various residues, and finally, these altered electrostatics of maspin-S176 and maspin-P176 rewired the polar contacts that abolished the allosteric control in protein. By combining these factors, the altered electrostatics can substantially affect the localization, and the preference of maspin binding partners, thus, culminating in a different maspin-protein (cofactor)-interaction landscape that could have been manifested with conflicting reports in previous studies. Here, the underlying reason has been highlighted and discussed that could be helpful for better therapeutic manipulations.

**Significance:** Protein altered functions in response to mutations are well documented, however, a slight perturbation in structure can lead to dramatic effects that are being felt at longer distances are rare. Here, we have reported that the substitution of Pro to Ser at 176^th^ position in Maspin can substantially alter the protein electrostatic interactions that can hamper the allosteric control. This could lead to different binding partners, localization preferences, and altered cytoplasmic retention duration resulting in functions that are not associated with normal protein. Moreover, the electrostatic attraction/repulsion can immensely affect the allosteric cohesion of protein resulting in unexpected outcomes.

## Introduction

Cancer growth can be prevented by multiple antitumor proteins that, if active during the early phase, trigger apoptosis and hinder the accumulation of transformed cells. Despite a larger number of antitumor proteins, the prevalence of cancer is growing and further studies are needed to rationalize the tumor initiation mechanism. Among the known factors for cancers including the inactivation of anti-tumor proteins, over-activation of pro-tumor proteins, and evasion of immune surveillance are notable. Once the tumor mass has been established, the cells invade and metastasize to other tissues, a process also termed epithelial-mesenchymal transition (EMT). This process has been executed by different transcription factors including Slug, Snail, Zeb1/2, and Twist, during embryogenesis and in malignant tumor cells in various combinations (Taube *et al*., 2010) (Yang and Weinberg, 2008; Schmalhofer, Brabletz and Brabletz, 2009; Micalizzi, Farabaugh and Ford, 2010). The EMT process can be characterized by loss of adherens junctions and relevant cell morphology changes from a polygonal/epithelial to a spindly/fibroblastic shape, increased motility, extracellular matrix degradation, and resistance to apoptosis. For instance, E-cadherin is the direct target of various transcription factors that could alleviate the suppression of motility and invasiveness (Peinado *et al*., 2004).

Among several tumor suppressor proteins, maspin is classified as a class II tumor suppressor that was first reported in 1994, and this protein prompts apoptosis, suppresses motility and metastatic related features (Zou *et al*., 1994)(Khalkhali-Ellis, 2006). Maspin is a non-inhibitory serpin (grouped with clade B serpins) owing to its short reactive cite loop (RCL) than those serpins who inhibit proteases, maspin cannot undergo a conformational change from stressed to relaxed conformation to inhibit protease activity. However, recent studies have furnished evidence for its functions in cellular attachment, motility, apoptosis, and angiogenesis that making it an attractive target for various forms of malignancies (Sheng *et al*., 1998; McGowen *et al*., 2000; Schaefer and Zhang, 2003; Berardi *et al*., 2013).

Maspin expression has been reported in various malignancies including gastric (Kim *et al*., 2005)(Zheng and Gong, 2017)(Tanaka *et al*., 2020), and its reactivation can suppress the breast tumor in xenograft models (Beltran *et al*., 2007). Maspin could be a biomarker for colorectal cancer (Banias *et al*., 2017), lung cancer (Berardi *et al*., 2015), oral cancer (Shpitzer *et al*., 2009), and cervical lesions (Manawapat-Klopfer *et al*., 2016). The maspin expression was evaluated in a wide range of cancers and found to be over/under-express depending upon the type of cancer. There are three coding region polymorphic forms of maspin has been reported, among these, P176S (rs2289519) has been linked to promoting gastric cancer (Jang *et al*., 2008), and V187L (rs2289520) has been associated with oral cancer (Yang *et al*., 2016). Maspin-P176 in conjunction with maspin-L187 decreases the risk of esophageal squamous cell carcinoma, while mapin-S176 (with maspin-V187) exerts the opposite effect (Meng *et al*., 2015). The homozygous maspin-S176 also has been shown to link with gallbladder cancer (Mahananda *et al*., 2021). However, how a minor variation in the non-structured region can drastically alter the function of the antitumor protein to pro-tumor protein is still a mystery.

Structural biology has advanced our understanding of the three-dimensional structures of various proteins, however, lacking the dynamic picture of these proteins hinders the functional aspect. The detailed analysis and resolution offered by molecular dynamic simulations (MDS) bridge the gap between structural biology and the dynamic nature of proteins (Hollingsworth and Dror, 2018). Here, in this study, we have analyzed the proteomics data from 80 early-onset gastric cancers and found out that maspin has been overexpressed in most of the samples. Furthermore, long MDS has been performed for two polymorphic forms of maspin-P176 and maspin-S176, and extensive energetics analysis was carried out. It has been shown that though, the residual contacts altered to a small scale, the per-residue and paired-residue energetics have been perturbed significantly. The rearrangement of the polar network and disturbance of allosteric signals are evident that make the maspin with small structural differences interact with different partners and result in opposing effects.

## Materials and method

### Maspin modeling

The three-dimensional coordinates for maspin protein have been downloaded from RCSB protein databank PDB ID: 1WZ9 (Law *et al*., 2005), and the missing loop residues (amino acids 335-345) have been built using the Swiss-Model online webserver (Waterhouse *et al*., 2018). After model building, the built model was compared against the 1XQG (Al-Ayyoubi, Gettins, and Volz, 2004) and the loop was in agreement with the crystal structure. For maspin-P176, the serine has been mutated in Chimera v1.1.3 (Pettersen *et al*., 2004) using Dunbrack 2010 (Shapovalov and Dunbrack, 2011) rotamer library with subsequent 500 steps of steepest descent energy minimization with 0.002 Å step size with an update of 100 conjugate gradient of same step size with none of the atoms fixed during minimization.

### Molecular Dynamic simulations

All molecular dynamics simulations (MDS) have been performed in GROMACS v2019.6 (Abraham *et al*., 2015) using AMBER99SB-ILDN forcefield (Lindorff-Larsen *et al*., 2010) with a dodecahedron box filled with rigid water model TIP3p water (∼20300) model (Jorgensen *et al*., 1983) and periodic boundary conditions were applied in all directions. The system has been neutralized and NaCl concentration was adjusted to 0.1 M (48 Na^+^, 41 Cl^-^). The system was energy minimized up to 500 kJ/mol/nm using the steepest descent algorithm and two-step equilibrations were performed. Initially, the temperature was adjusted to 310 K for 1 ns using the V-Rescale method (Bussi, Donadio and Parrinello, 2007) with 0.1 ps time constant, and then isotropic pressure (1 bar and 2.0 ps time constant) was applied to the system following Parrinello-Rahman algorithm (Parrinello and Rahman, 1981). During equilibration, the position restraints were applied to avoid any structural distortion. Particle Mesh Ewald (Darden, York, and Pedersen, 1993) has been used to treat long-range electrostatic interaction with cubic interpolation, and a 10 Å cutoff was used for short-range Coulomb and Van der Waals interaction. For each system, five independent production simulations have been performed for a length of 1 μs each starting from a different random seed with only hydrogen bonds constrained using LINCS (Hess *et al*., 1997) and a trajectory snapshot has been saved with 10ps. Therefore, a total of 10μs simulation has been conducted and, most of the analysis has been conducted for the last 500 ns with a 100 ps gap making a total of 25000 snapshots for each analysis.

### Nonbonded Interaction Energies calculation

The residual and paired-residue non-bonded interaction energies have been calculated between maspin-P176 and maspin-S176 as follows; the differential residual average non-bonded energy of residue i is:

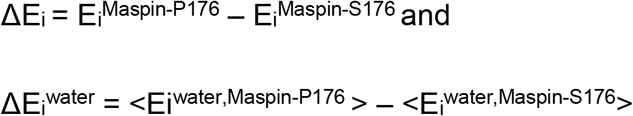

The paired-residue interaction energy for residue i and j in maspin-P176 and maspin-S176 has been calculated as:

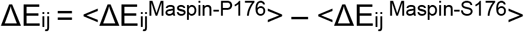

The symbols <> indicate the ensemble average for the respective energy terms (electrostatic (E^Elect^) and Van der Waals (Vdw, E^Vdw^)), however, the Vdw energy terms are negligible than electrostatic interactions, therefore, primarily E^Elect^ has been discussed in details.

### Differential Contact Map

The contact between two residues has been defined as if the distance between heavy atoms was less than 4.5 Å with four residues apart (Yuan, Chen, and Kihara, 2012). The contacts have been calculated for the last 500 ns of each trajectory (total frames of 25000), and the interatomic contact fraction is defined as: f_ij_ = n_ij_ / N; where n_ij_ is the number of frames contact between atom_i and atom_j is present, while N is the total number of frames. For each residue, the individual contacts have been summed, thus a value of more than one is possible.

### Allosteric coupling intensity analysis

The allostery among the residues has been calculated using Ohm which employs the concept of perturbation propagation to determine the allostery in proteins (Wang *et al*., 2020). As the allosteric communications depend on the interatomic contacts, thus, a contact distance of 4.5 Å has been used, with a probability of perturbation from one residue to another set to 0.05 and the number of perturbation round was set to 10000. The value of α was set to 0.5 to capture the possible allosteric probability. The remaining parameters were kept to default values as provided by the algorithm.

### Analysis

Most of the analyses have been performed using tools provided with GROMACS v2019.6 and the amber trajectory processing tool (cpptraj) (Roe and Cheatham, 2013). The 3D protein figures are generated using Chimera v1.1.3, and plots are generated using MS Excel, and Matplotlib library.

### Statistical test

For the statistical test, the data were binned into 25 sets, and an average of each set was then used to assess the significance (p-value < 0.05) using a two-tailed student T-test in SciPy with default parameters (Virtanen *et al*., 2020).

## Results

### Maspin was stable during MD simulation

From the gastric cancer dataset of 80 patients, the maspin has been identified to be upregulated in 55 (69%), and downregulated in 8 (10%), while it was not detected/significant in 17 (21%) of samples (Supplementary method, Supplementary Fig. 1). Maspin has been attributed as the antitumor protein where it prevents metastasis, a hallmark of cancer, and functional loss of this protein can enhance tumor potential (Zhang *et al*., 2000). Normally, maspin-P176 functions as a tumor suppressor, however, the substitution of P176S can alter the functions making it a protumor protein, prompting this study to reveal the functional aspect of these polymorphic forms (Fig. 1A). From modeling, both of these forms are nearly identical (RMSD between Cα 0.16 Å over 375 atom pairs), however, when the inter-residue contacts were analyzed, substantial differences were noted (Fig. 1B) that indicate the sidechain rearrangements in maspin-S176 and maspin-P176. Therefore, to further study the impact of side-chain rearrangement, each polymorphic form *i.e*. maspin-P176 and maspin-S176 has been subjected to five independent 1μs long all-atom molecular dynamics simulations.

**Fig 1.**
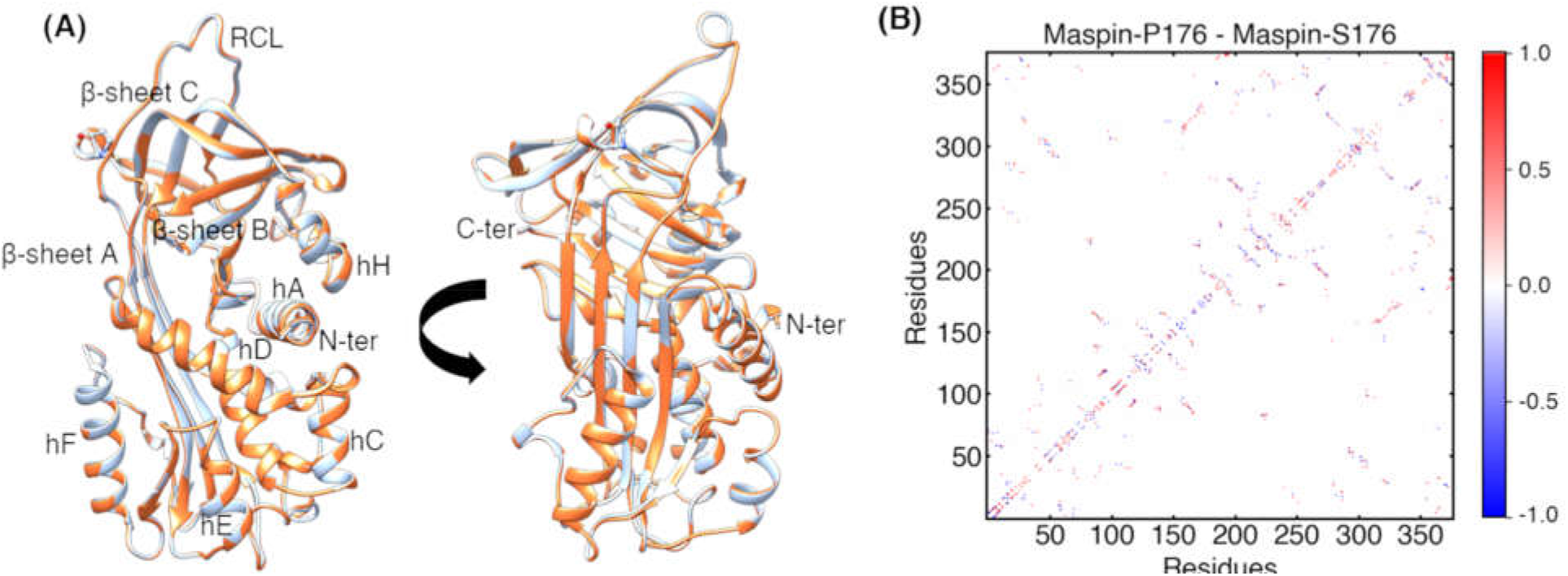
Structure and contact analysis of Maspin. (A) 3D structure of maspin (P176 in light blue and S176 is in brown). The prominent secondary structural elements are also labelled. (B) Differential contact map analysis of maspin-P176 and maspin-S176 from the energy minimized structures. C/N-ter, C/N terminal, RCL, reactive center loop.

As the polymorphic forms differ in single amino acid (Ser or Pro at 176) on a loop region, none of the proteins underwent any conformational change (Supplementary Fig. 2). After the initial adjustment for around ∼300 ns (the RMSD change was ∼0.25 nm), the root-mean-square deviation (RMSD) of each protein remained stable for the remaining length of simulation (∼0.1 nm) (Supplementary Fig. 2A). Similarly, for root-mean square fluctuation (RMSF), most of the residues fluctuate proportional to each other, however, the non-structured region is showing higher fluctuation such as residues 61-83 (loop between hC-hD and hD); 102-113 (hE and proceeding loop), the loop between hF-s3A (138-150), the loop between s3B-hG (234-242), and the prominent RCL region 334-345 (Supplementary Fig. 2B). The other structural measures such as solvent accessible surface area (SASA) were also consistent (Supplementary Fig. 2C).

The average number of intra-protein hydrogen bonds was higher for maspin-S176 (290.4 vs maspin-P176 286.4), however, the total number of hydrogen bonds was substantially lower than that of crystal structure (*i.e*. 412) (Law *et al*., 2005). The hydrogen bonds between protein and water were not significantly different (two-tailed p-value = 0.6) (Supplementary Fig. S2D, E) indicating that the internal reordering of the bonding network is plausible.

The water count in the first and second shell around the maspin-P176 (1379.3±25 and 2200±34.5) and maspin-S176 (1370.9±26, and 2191.1±38.3 respectively) are in close range. Similarly, the fraction of native contacts (Q) indicated that the proteins are at an intermediate-active state with preserving ∼0.43 of native contacts in both cases (Supplementary Fig. 2F) (Giri Rao and Gosavi, 2018). As it has been shown that these proteins can fold to meta-stable conformation before folding into stable conformation, which assists in their functions. Moreover, the maspin-S176 showed a lower packing density with a higher number of cavities signifies that Mapin-S176 is less compact (Supplementary Fig. 3).

Recently, maspin residues from 87-114 were proposed to harbor nuclear localization signal, and their exposure impact maspin localization (Reina *et al*., 2019). When the exposure has been calculated for the entire length, maspin-S176 showed slightly higher solvent accessibility than maspin-P176, however, for 87-94 (s2A), maspin-S176 was significantly higher referring to the higher possibility of this variant being nuclear-localized (Supplementary Fig. 4).

### The differential contact map highlighted the pivotal difference in P176S

As maspin in either form is stable, and it has already been shown that the side chain contacts could be altered, the residue contact map has been analyzed next. The residue communication occurs through residual contacts among themselves that exert influence over the neighboring residues. Due to the dynamic nature of proteins, these contacts are dynamically formed and break over time. This can provide an intuitive way to analyze the close collaboration and allosteric influence of various residues. In Fig. 2, the blue and red dots represent the higher fraction of contacts in maspin-S176 and maspin-P176 respectively, with a maximum of ∼30% change in contact frequency has been observed (Supplementary Table 1). The change in contacts is in close vicinity referring to the rewiring of contacts and allosteric communications. For instance, the difference in contacts between residues S40-F70 (−27.2, hB-hD, can influence collagen-binding), K268-H344 (−24.8, influence RCL mobility), Y84-K170 (−23.1, help expose Y84 and phosphorylation), M172-S333 (−19.7, vicinity of mutated site and base of RCL) and Y84-F167 (−18.3, influence hydrophobic packing) were increased in maspin-S176, whereas, Y84-H225 (29.9, protection of shutter region), E218-F229 (28.9, β-sheet B, core hydrophobic region), F60-F70 (23.6 hC and hD, may alter collagen-binding), E289-K294 (19.8, solvent-accessible), and T241-P353 (17.1, influence hG movement) were decreased in maspin-S176. Similarly, the base of RCL has shown substantial contacts with the region around 176 in maspin-S176 highlighting the mobile side-chain interactions.

**Fig 2.**
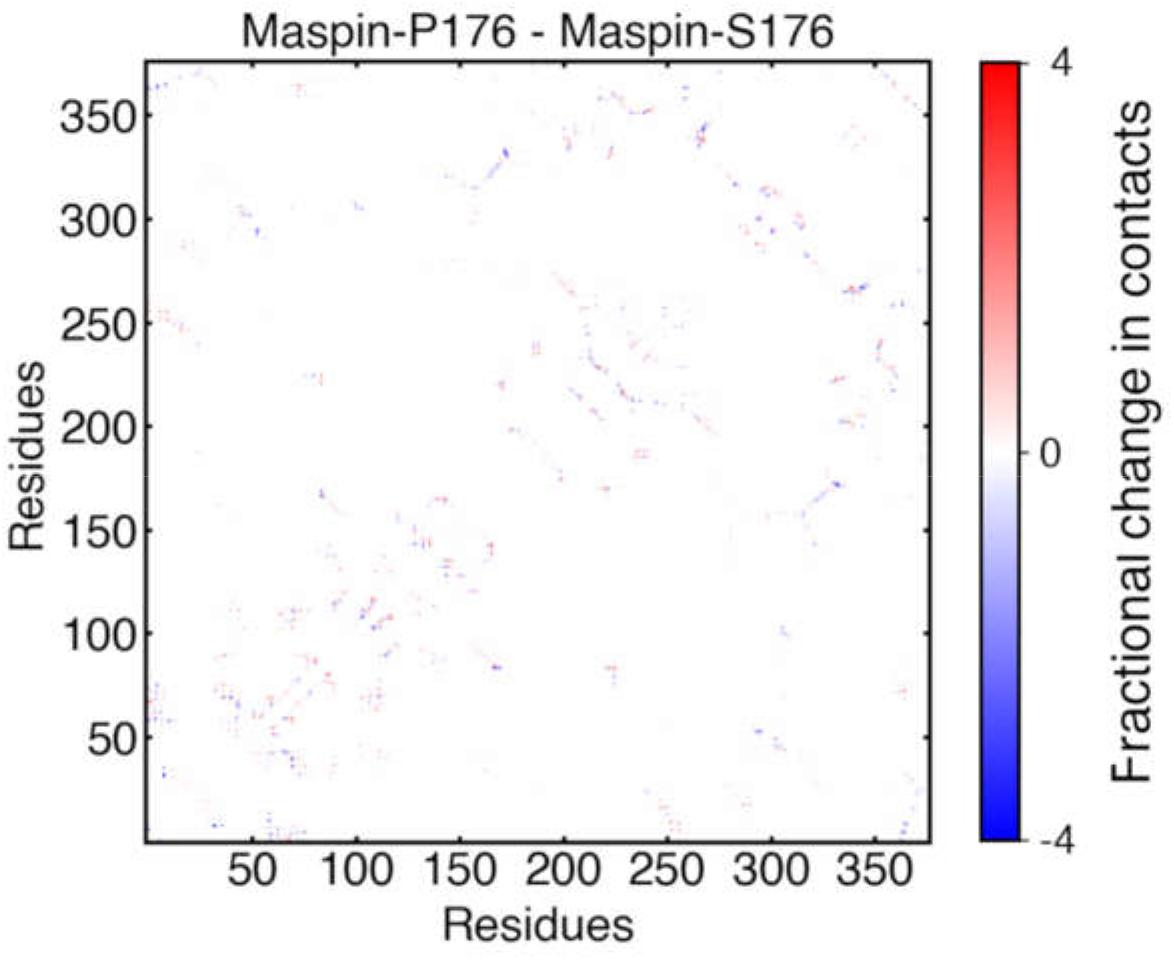
Differential contact frequency map. The contacts among heavy atoms only have been calculated with cutoff of 4.5 Å and contacts measured for only residues with 4 residues apart. The maximum fractional change was found to be 6.4, however, for matrix representation, the color was anchored at 4 for better visualization, as only 0.39% contacts are beyond this range.

### Rewiring of Electrostatic interactions is evident in P176S

As it has been evident that there are subtle changes in inter-residue contacts, and all the non-bonded interaction arises from these contacts, therefore, it is reasonable to argue that the nonbonded interactions have also been perturbed (Fig. 3). The non-bonded interaction energies (electrostatic (E^Elect^) and Van del Waals (E^Vdw^)) have been calculated for each residue and between protein and water. The total non-bonded interaction energy (E^Total^ = E^Elect^ + E^Vdw^ and ‹ΔE^Total^› = ‹Maspin-P176 ^Elect^+^Vdw^› - ‹Maspin-S176^Elect^+^Vdw^›) is heavily influenced by the change in electrostatic energy, whereas the ΔE^Vdw^ is nominal (mostly < 2 kcal/mol). As the ΔE^Total^ is proportional to ΔE^Elect^, thus, it is rational to focus on ΔE^Elect^.

**Figure 3.**
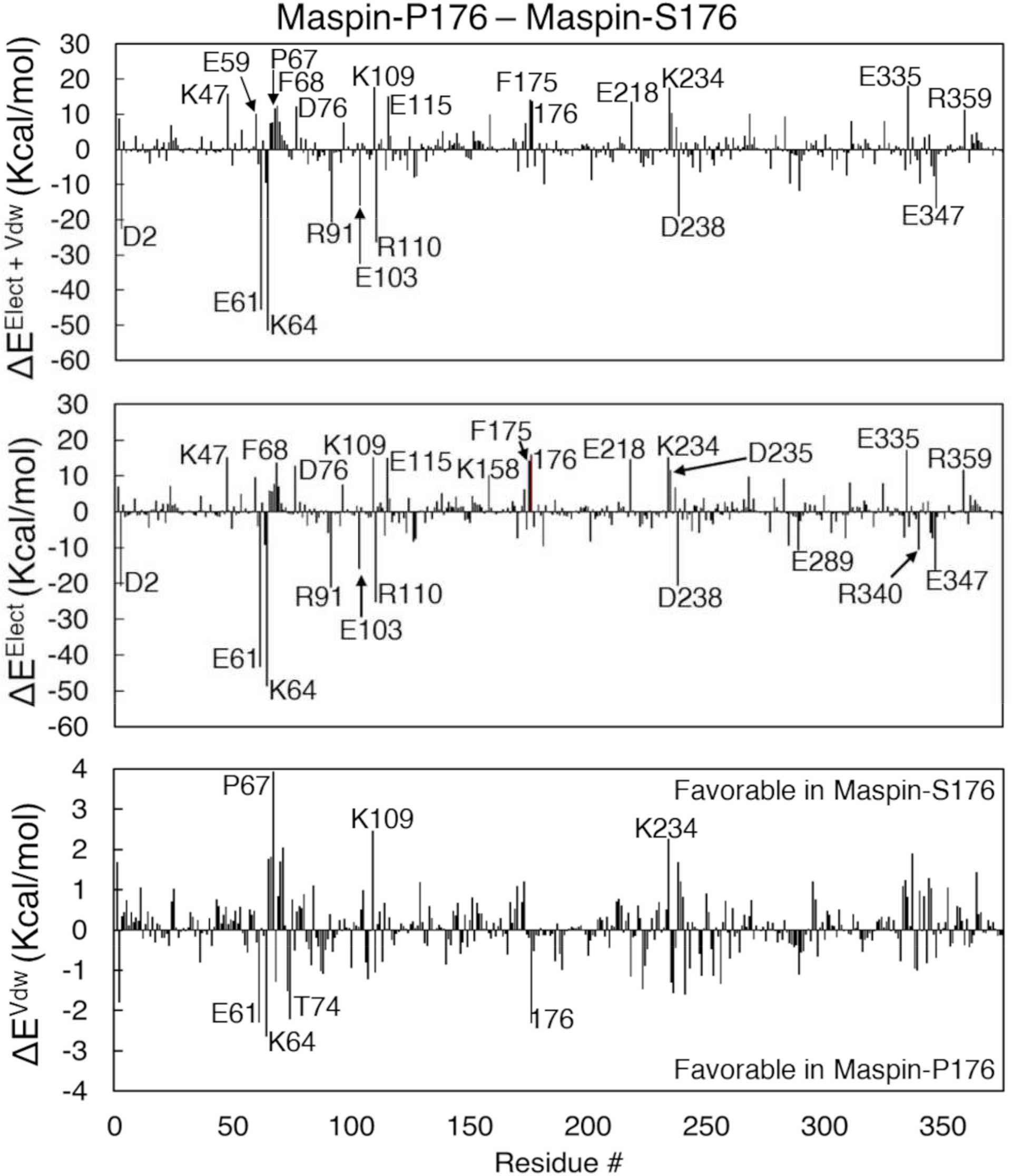
Per-residue energy perturbation in Maspin. The average energy change has been calculated from 5000 snapshots of each trajectory. The total interaction energy is the sum of nonbonded electrostatic and Van der Waals energy terms.

The residues with significant perturbed electrostatic energy are scattered across the protein, including the mutated position 176 (15.98 kcal/mol). In the figure, the positive and negative values indicate favorable and unfavorable ΔE^Elect^ energies respectively in maspin-S176. Among these residues, E61 (−43.2 kcal/mol) and K64 (−48.7 kcal/mol) showed the highest unfavorable energy for maspin-S176 and are present on the hC-hD loop, far from the mutation site. Moreover, E335, E340 and E347 are present on/around the RCL, whereas, E335 being closer to 176^th^, could participate in altering the dynamics of RCL. R91, K109, and R110 are part of positive residue clusters that severely altered their ΔE^Elect^, which is known to bind cofactor/proteins in many maspins (Sheng *et al*., 1996; Parker *et al*., 1999; Reina *et al*., 2019), where only R91 having unfavorable electrostatic energy in maspin-S176. ΔE^Elect^ in this region is also critical as it has been reported to act as nuclear localization signal and altered electrostatic could mask or unmask this region (Reina *et al*., 2019). E218 and R359 are at β-sheet B which could significantly alter the inter-residue distance and compactness of the protein (Supplementary Fig. 3). Similarly, an overall distance between acidic (D and E) and basic residues (K, R, and H) has been significantly higher in maspin-S176 (p-value < 0.01, Supplementary Fig. 5) that shows the electrostatic energies can alter the protein’s physical state. Concerning ΔE^Vdw^ interaction energies, P67, K109, and K234 showed favorable, while E61, K64, T74, and S176 were unfavorable ΔE^Vdw^ in maspin-S176.

The residual interaction energy between protein residues and water is also dominated by ΔE^Elect^. Most of the residues are polar and are showing favorable ΔE^Elect^ in maspin-S176 with water, which is rational and provides solvent-mediated dipole orientation to fine-tune the protein solubility and functions (Qiao *et al*., 2019). In the case of ΔE^Vdw^ energy, only Y112 and S176 are prominent (Supplementary Fig. 6). The Y112 with phenol side chain can be phosphorylated resulting in altered functions.

### Propagation of electrostatic perturbation in P176S maspin

As it is now known that the mutation causes significant changes in the electrostatic interaction, it is good to know how these perturbations travel across the protein. For this, the pairwise residue interaction (ΔE_ij_^Elect^ = <E_ij_>^P176^ − <E_ij_>^S176^ has been calculated for all the interacting pairs (Fig. 4, Supplementary Fig. 7, the numerical values of | ΔE_ij_ | > 4 kcal/mol are given in supplementary Table 2). The interacting residues have been drawn as a network to visualize how the signal travels from the mutated site (Fig. 5).

**Figure 4.**
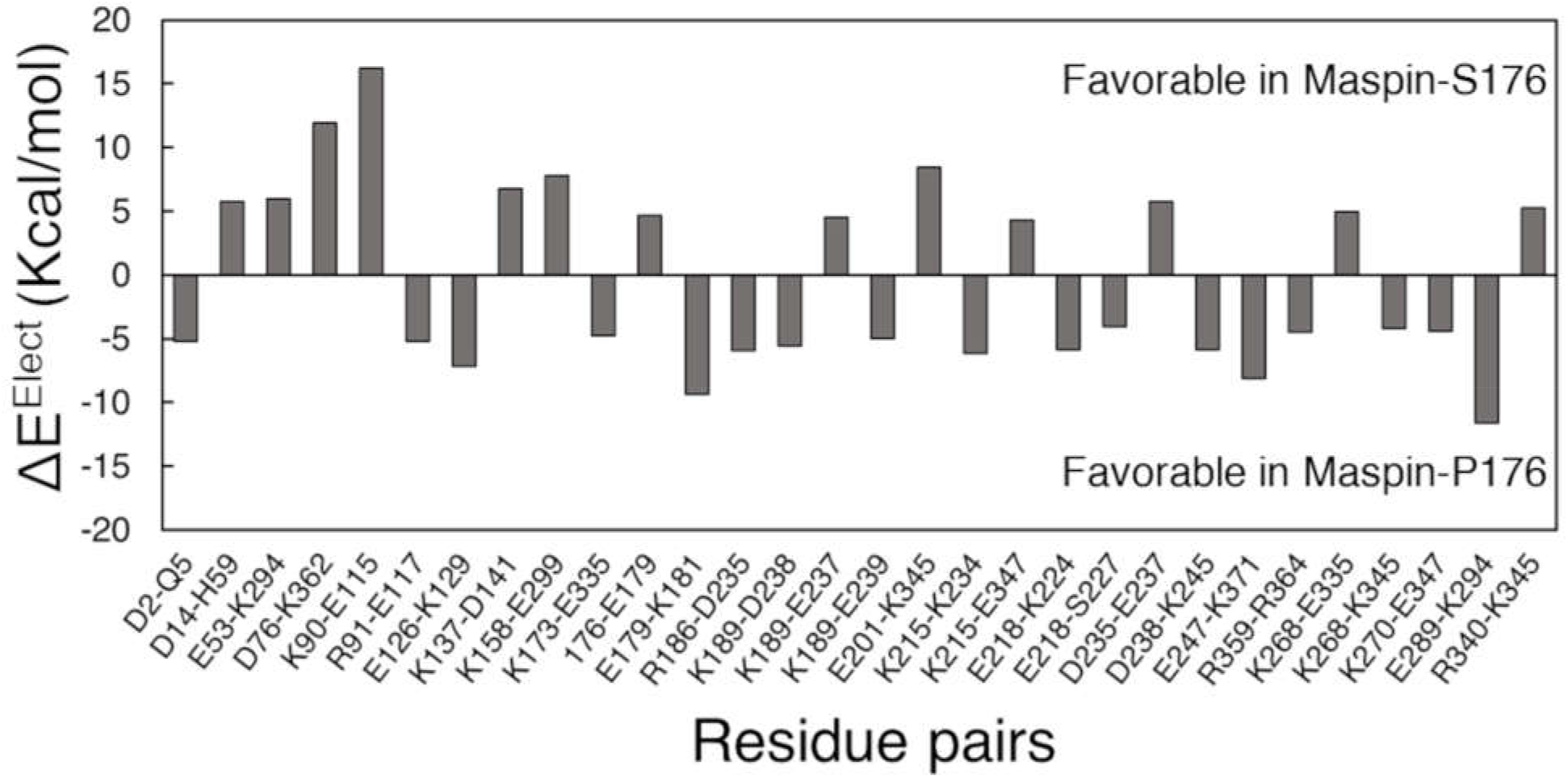
Pair residue ΔE^Elect^ interaction energy. The ΔE^Elect^ is substantial between P176 and S176 (only residue pair with >±4 Kcal/mol are shown).

**Figure 5.**
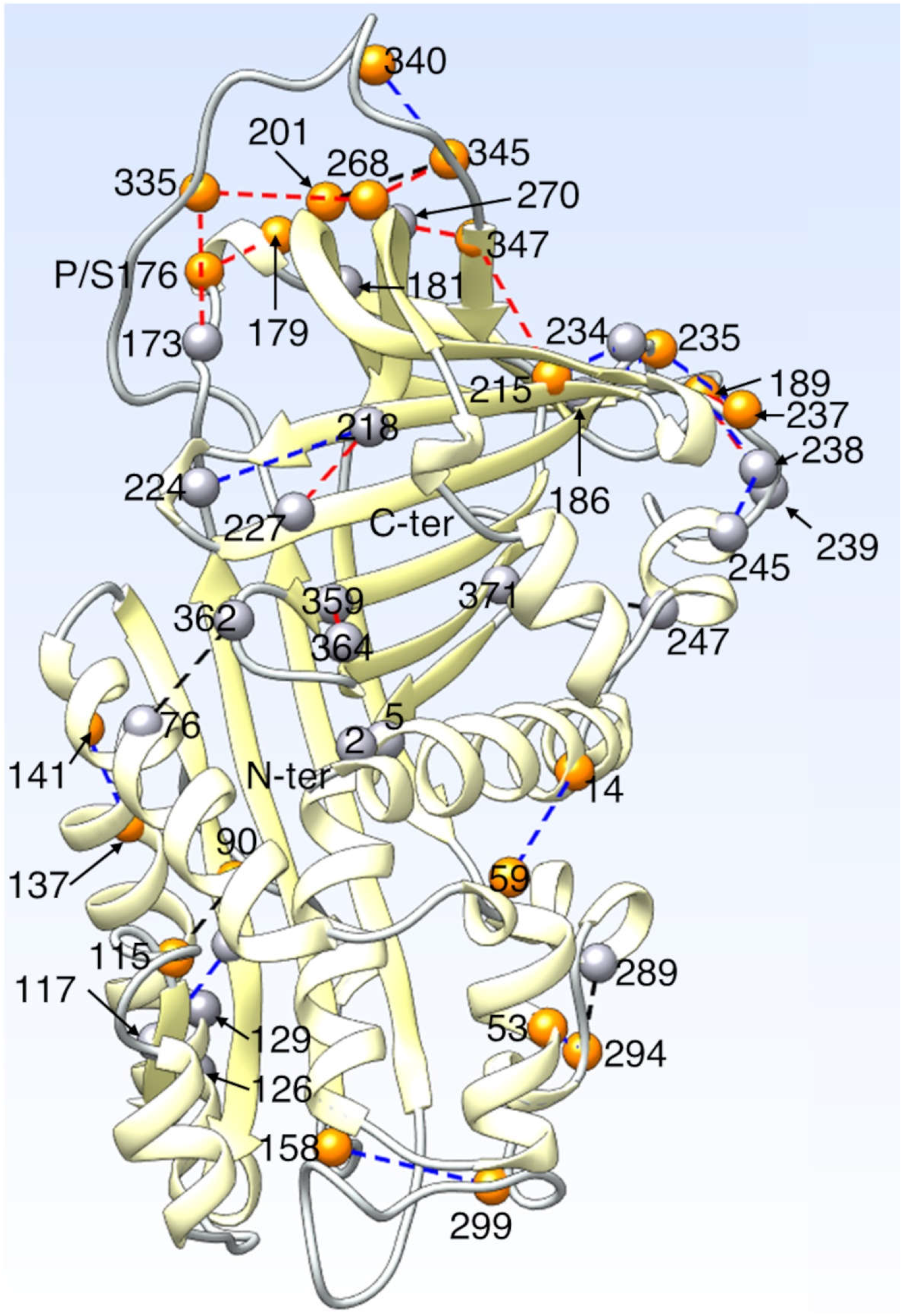
Network view of paired-residue ΔE^Elect^ interaction energy with. only residue pair with >±4 Kcal/mol are shown. The black dotted lines represent | ΔE^Elect^ | > 10 kcal/mol, blue denotes the |ΔE^Elect^| > 6 kcal/mol, and red denotes | ΔE^Elect^ | > 4 kcal/mol. The residues with favorable interactions in Maspin-S176 are in orange, while with unfavorable interactions are in gray.

The paired residue-wise electrostatic interaction energy is substantially different for various residue pairs. For instance, D14-H59 (5.7 kcal/mol) and K90-E115 (16.2 kcal/mol) form salt bridges, and it can be shown that both are showing favorable electrostatic interaction in maspin-S176. These pairs are far from the mutation site yet they display substantial electrostatic influence. Moreover, the salt-bridge formation between K90-E115 and D14-H59 is energetically expansive and generally destabilizes the protein structure, however, such interactions can play role in determining specificity in various protein-mediated interactions (Hendsch and Tidor, 1994; Nakamura, 1996; Zhou and Pang, 2018). Similarly, E201-K345 (8.4 kcal/mol) displayed a favorable interaction that might assist the RCL terminal region to get the anchor. The effect of the mutation was unfavorable in the vicinity of mutation, E179-K181 (−9.3 kcal/mol), where a polar substitution (P176S) might create charge repulsion. Similarly, E289-K294 (−11.6 kcal/mol) and E247-K371 (−8.1 kcal/mol) also exhibit the unfavorable interactions.

As can be seen, the mutation from Pro to Ser causes a local electrostatic disruption that triggers the neighboring residues, such as 173, 179 and E201 to be perturbed. The 173 has an interaction with 335 (the base of RCL) that then propagate the signal through β-sheet C and RCL to the stretch of residues ranging from K234, D235, E237 D238, and E239 that can alter the electrostatic potential (Fig. 5). Similarly, hG is also close to these residues that has been reported to move resulting in an open conformation in crystallographic structure. This hG has been implicated in collagen binding and altering the overall kinetics of protein-protein interaction and acting as a conformational switch (Law *et al*., 2005). Moreover, E218 is in the s2B that could significantly change the inner core potential and can alter the interaction between the T259-R364 residue pair, which is in direct connection to hA. From the β-sheet A, there are only a few residues such as K90, R91, D115, and E117 that are affected indicating the inter-residue ΔE^Elect^ alteration can propagate the signal to a longer distance.

Concerning Vdw interaction energy, only a small stretch of protein exhibits drastic changes, such as, D351-H352 was highly stable in maspin-S176 while H352-P353 was stable in maspin-P176 (Supplementary Fig. 7). These residue positions are in close proximity to the mutation site that could feel the polar or nonpolar energetic differences the highest. Similarly, H360 substantially gained stability with T363 and D76. Moreover, N163-H320 are buried in the core which is also more stable in maspin-S176.

### P176S substitution alters the ionic bonding network

As the substitution of P176S causes significant alteration in electrostatic interactions, and almost all of these residues are polar, they are capable of forming ionic bonds (salt-bridges and hydrogen bonds) with each other. Consequently, it is intuitive to evaluate the frequency of ionic interactions between residues. In figure 6, the representative polar contacts have been shown (Fig. 6). It should be noted that these contacts are dynamic, and can be in different states in different conformations.

**Fig 6.**
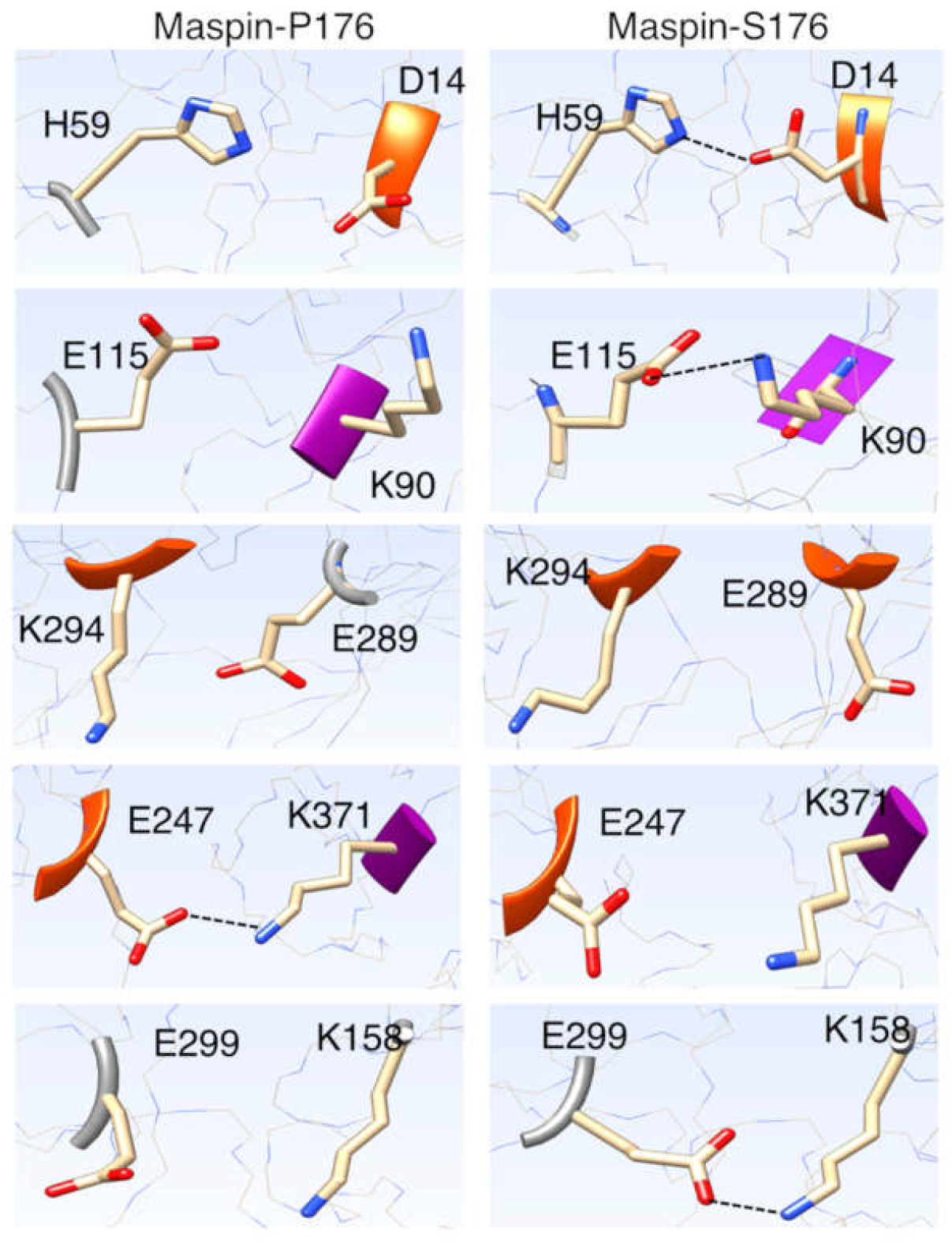
Representative snapshots of residues with polar contacts. These contacts are dynamic, and represent ensemble average. The contacting residues are in stick representation with heteroatomic colored and hydrogen are hidden for clarity. The secondary structure is colored (α-helix is in orange, β-strand is in purple, loop is in gray). The remaining protein is in line representation.

The probability distribution of the minimum distance of side chains and the differential hydrogen bond (HB) frequencies (Hbond^P176^ – Hbond^S176^) have been given in figure 7. For most of the pairs, the overall pattern is similar albeit the intensity varies so is the HB frequencies (negative values indicate a higher presence of HB in maspin-S176). For instance, the residue pairs K90-E115, D14-H59, E247-K371, E289-K294, and K158-E299 all exhibit a strong single peak at 0.2 nm that indicates ionic interaction (salt bridge or hydrogen bond). The K90-E115, D14-H59, and K158-E299 are more stable in maspin-S176, while, E247-K371 (hG and s5B) and E289-K294 (hH and proceeding loop) are unstable in maspin-S176. In the case of D76-K362, E201-K345, and D238-K245 all are showing two or more peaks with D76-K362 and E201-K345 being more stable in maspin-S176. There is a single pair that had a non-overlapping population shift, D235-E237 with maspin-P176 having stable interaction and higher hydrogen bond propensity than that of Mapisn-S176. Finally, there is no hydrogen bond was found between R340-K345 mainly due to the disordered nature of RCL. These population shifts of polar ionic bonds can change the inter-residue distance, electrostatic interactions, allosteric signaling and communication, and the continuity of charges that could significantly modulate protein binding mode.

**Figure 7.**
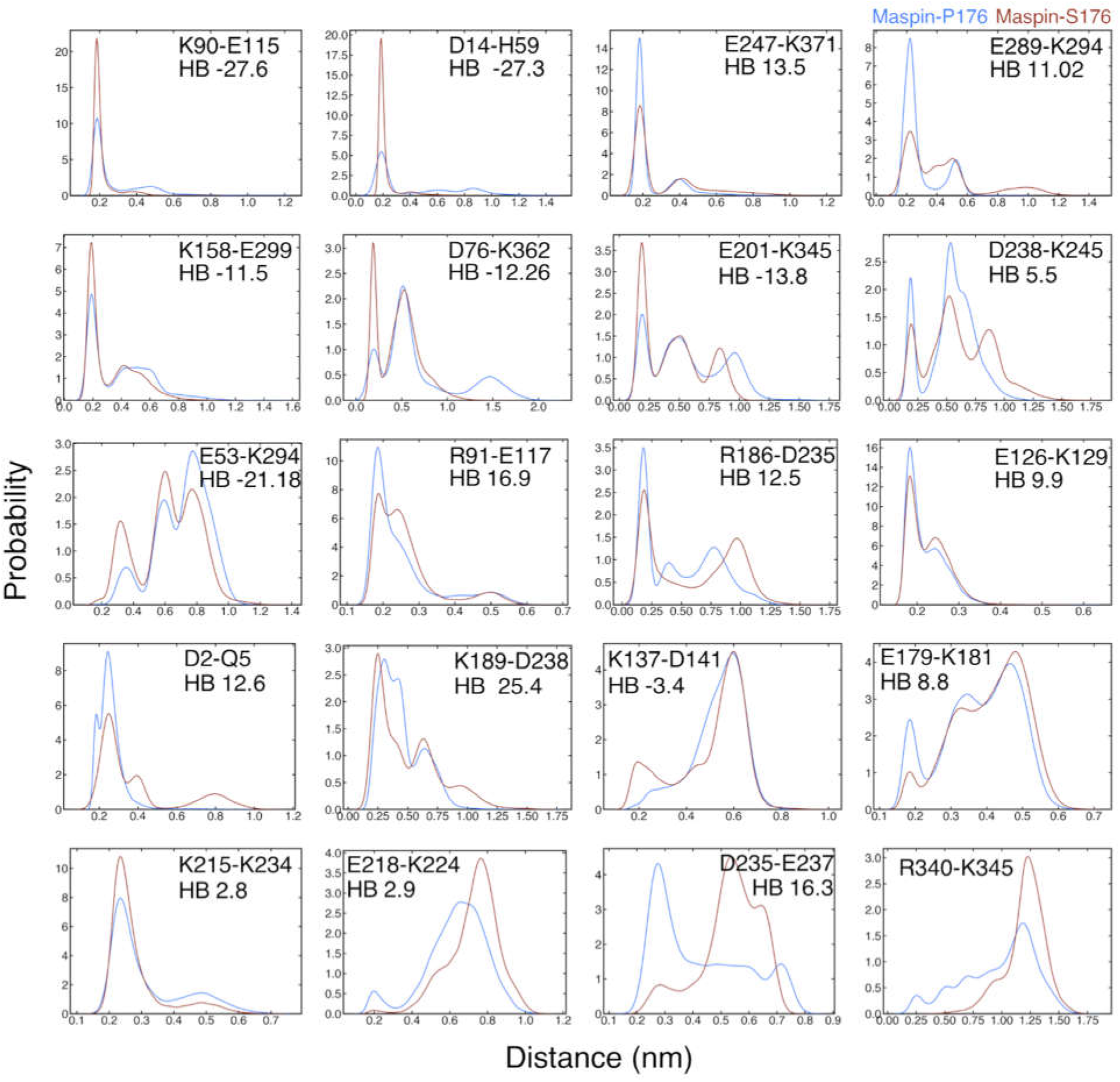
Paired residue distance distribution and hydrogen bond occupancy. The minimum distance between residue pair has been calculated for the last 5000 snapshots of each trajectory and plotted as the distribution graph for only pairs with | ΔE^Elect^ | > 5 Kcal/mol. The change in hydrogen bond (HB) occupancy (Maspin-P176^HB^ – Maspin-S176^HB^) has been given as well (except R340-K345 as there is only single HB was observed). The presence of polar contacts such as salt-bridge or HB can be analyzed by the peak at 0.2 nm.

The salt-bridge formation between K90-E115 and D14-H59 has been reported in the crystal structure that carried Ser at 176 (Law *et al*., 2005), however, in the case of P176, these salt bridges dramatically reduced. This indicates that the electrostatic interaction can have a ripple effect. Moreover, it can be inferred that the stronger polar interaction between these residues should stabilize the protein as has already been reported for various thermophiles (Kumar and Nussinov, 2001).

### P176S causes the loss of allosteric coordination in maspin

The mutation caused the electrostatic interaction and bonding network to shift dramatically, the next question may be whether these rearrangements can have any impact on allosteric interaction within protein? As the allostery is pivotal in protein functioning where any improper signaling can heavily influence the functional capability.

The mutation P176S caused a dramatic decrease in allosteric coupling intensity (ACI, a measure to show the allosteric influence from one residue to another) (Fig. 8A, numerical values with | ΔACI | > 600 are provided in Supplementary Table 3) (Wang *et al*., 2020). In the case of maspin-S176, almost all the residues have shown a substantial drop in ACI values. The reduction is prominent for residues 175, 185, 186, 197, 198, 204, 206, 271, 273, and 353 where the ΔACI >200. Most of these residues are in the vicinity of mutation. The pathways followed by the signal propagation have a lower weight in maspin-S176 in comparison to maspin-P176 and the paths are mostly similar (Fig 8B, C). The signaling path is only divergent from 360, in maspin-P176, it mostly goes to 363 while in maspin-S176, 365 is the prominent candidate. From 363/365, different routes are being followed to reach the target residue (K64).

**Figure 8.**
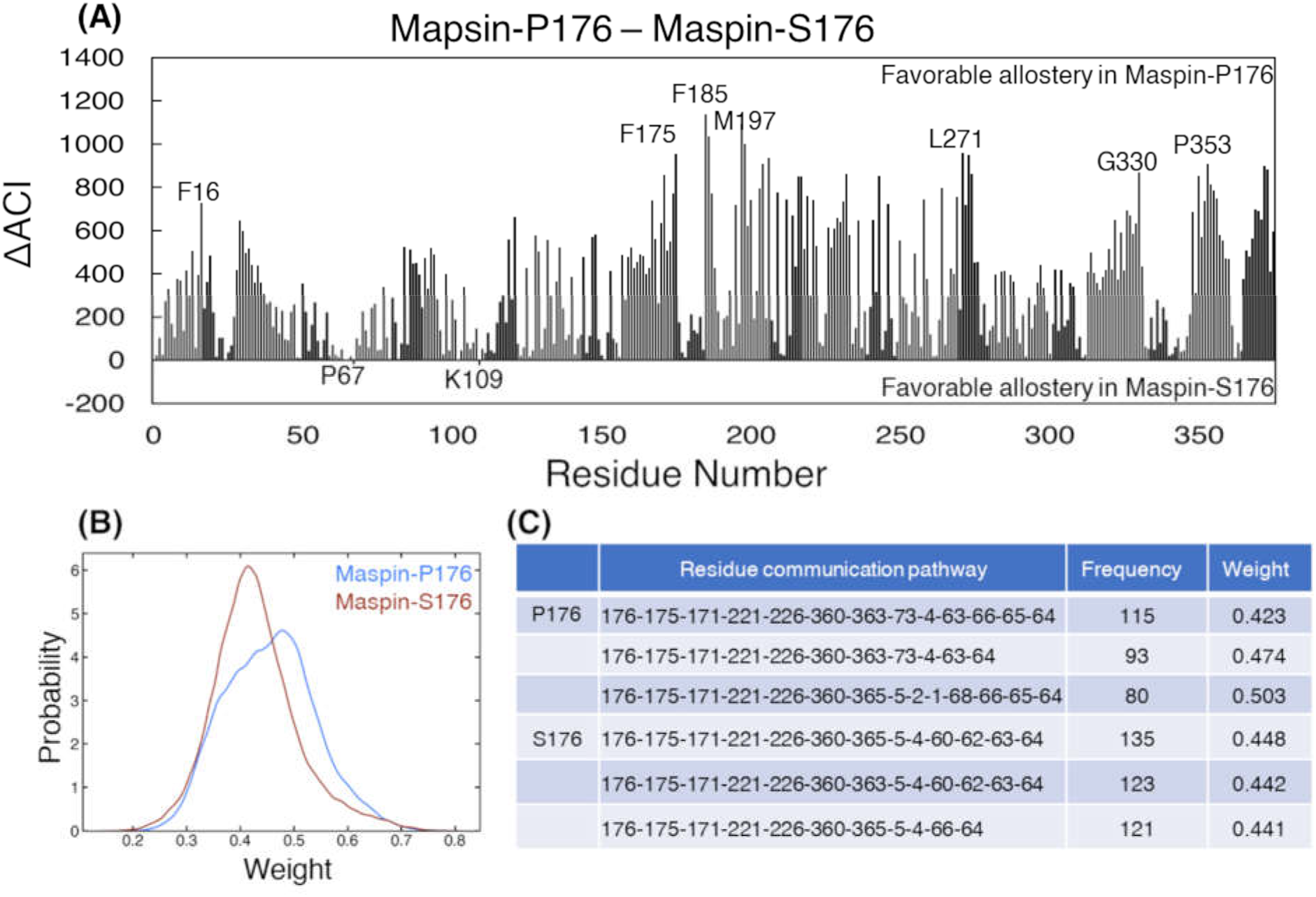
Allosteric coupling intensity (ACI). The ACI values have been calculated using perturbation propagation algorithm that analyzes the probability of propagating the signal from allosteric site (in this case, 176^th^ residue) to the target site (here 64 chosen as the active residue as it shows the highest ΔE^Elect^). The residues with positive values represent decrease, while residues with negative values represent increase in ACI in Maspin-S176. Few residues have been labelled for reference. **(B)** The distribution of weights from highest probable paths from allosteric site to target site using 25000 snapshots (Kolmogorov-Smirnov p-value << 0.001). **(C)** The top 3 pathways have been given for each protein along with the frequency of frames.

## Discussion

The present study aimed to highlight the critical elements that govern the differential role of maspin in structural and dynamical contexts using long unbiased MDS. In MDS, the non-inhibitory maspin is stable and in the active state conformation that exhibits an inter-residue and paired-residue electrostatic perturbation in response to mutation. These electrostatic perturbations have reduced the allosteric coupling within the protein that could have altered the whole maspin-interaction landscape resulting in altered interacting partner preference (Zhang, Witham, and Alexov, 2011), cellular localization and stability (Strickler *et al*., 2006; Kudriaeva *et al*., 2019) and posttranslational modification propensity (as reviewed in (Bianchi *et al*., 2020; Vascon *et al*., 2020)). To further explore the protein-protein interactions (molecular docking simulations) that consider the significant changes in electrostatic interactions required further studies.

The localization and cellular environment of maspin largely dictate its pro- or antitumor functions (Bernardo *et al*., 2015; Dzinic *et al*., 2017; Sakabe *et al*., 2021)(Zhang *et al*., 2000; Ngamkitidechakul *et al*., 2003; Dzinic *et al*., 2013). Recently, it has been reported that maspin can translocate to the nucleus through active and passive mechanisms, and it also harbors nuclear localization signals, spanning from 87 to 114 (s2A and hE) (Reina *et al*., 2019). This region holds a strong salt bridge, K90-E115, in maspin that could help to mask/unmask this region and regulate nuclear localization. In our analysis, maspin-S176 is slightly more exposed (Supplementary Fig. 4A & B) that should favor nuclear localization, however, for antitumor activity, there may be more than one variable involved. Further, maspin cellular localization is regulated by cellular confluency and epidermal growth factor receptor-related pathways implicating their role in maspin functions (Longhi *et al*., 2021). Nuclear localized maspin has been reported to inhibit breast cancer proliferation (Tanaka *et al*., 2020). Similarly, Dzinic et al, 2013 have reported that nuclear-localized maspin effectively inhibits HDAC1 functions and suppresses the growth of multiple cancer cell lines including lung and prostate, and nuclear localization is favored by mutation around the RCL region (maspin-D346E) (Dzinic *et al*., 2013). Both of these residues are negatively charged and should not alter the isoelectric point; however, a longer side chain of glutamate could offer more interacting opportunities.

Maspin is known to interact with many proteins including interferon regulatory factor 6 (Bailey *et al*., 2005), collagen (type I and III) (Blacque and Worrall, 2002), urokinase-type plasminogen activator and its receptor (Pemberton *et al*., 1997), β1-integrin (Cella *et al*., 2006), heparin (Sheng *et al*., 1996), heat shock protein 70 and 90, glutathione *S*-transferase (Yin *et al*., 2005), and Bax, an apoptotic related protein (Liu *et al*., 2004). Similarly, the electrostatic interactions are critical in ternary complex formation in transcriptional adaptor zinc-binding domain 1 protein illustrating vital electrostatic control in protein binding and localization (Wang and Brooks III, 2020). Further, in maspin-S176, the favorable ΔE_ij_^Elect^ interaction increased for surface residues that could modulate solvent interaction and hence stability and altered the binding partner preference (Fig. 5, Supplementary Table 2) (Strickler *et al*., 2006). Furthermore, the altered charge distribution in proteins can attract new partners such as in fatty acid-binding protein 5 (FABP5), ligand binding forces the basic residues to align and create a docking site for importin binding that subsequently translocate FABP5 to the nucleus (Armstrong *et al*., 2014). In maspin, a similar alignment can be achieved using altered inter-residue interaction (Reina *et al*., 2019).

In maspin, residues from hD, hE, and hG form a negative patch that binds with collagen, an integral cellular component and critical for angiogenesis, sharing a similar binding interface to other serpin family members such as pigment epithelium-derived factor (Blacque and Worrall, 2002; Meyer, Notari and Becerra, 2002; Law *et al*., 2005). Among these residues, E61 and D238 exhibited the unfavorable ΔE^Elect^ and hG harboring electrostatically repulsive pairs that could substantially impact the collagen-binding (Fig. 5, Fig. 6). Similarly, RCL has been reported for cell adhesion (Ngamkitidechakul *et al*., 2003; Dzinic *et al*., 2013), and mutation in this region may hamper its activity. Owing to its solvent-exposed and disordered nature, there were few residues whose energetics were significantly perturbed, such as E335, R340, and possibly E347. The mutation of R340 has been suggestive of loss of cell adhesion, and R340 is exhibiting unfavorable ΔE^Elect^ whose energy is being partly compensated by interacting with K345 in maspin-S176. Therefore, the altered electrostatics could hamper interaction with other proteins/cofactors (Fig. 5, Fig. 6).

Allosteric regulations are known to regulate a variety of protein functions such as protein binding (Lin *et al*., 1998), and complex formation (Soisson *et al*., 1997), and SNPs are known to affect the allostery (Tee, Guarnera and Berezovsky, 2019), allosteric polymorphism. The role of these mutations has been extensively studied for tailored medicine (Nussinov and Tsai, 2015; Nussinov *et al*., 2019), however, classifying P176S as a cancer driver required further studies.

In conclusion, the P176S mutation in maspin has altered the residual electrostatic energetics as well as created an altered paired-residue interaction pattern. These altered electrostatic charges can substantially change the binding partner preference and mode, localization, and stability that could influence the overall functions of the protein the in cellular environment.

## Supporting information

Suppplementary material

## Acknowledgments

This study has been funded by the National Research Foundation of Korea grants funded by the Korean Government (MSIT) (NRF-2015R1A5A1009024, NRF-2019M3A9D5A01102794).

This research was supported by the GRRC program of Gyeonggi province [GRRC-Kyung Hee 2018 (B03) to KPK].

## Authors contribution

MAA and KPK designed the study, KPK arranged the materials, MH, SC conducted simulations, and MAA and KPK analyzed the results and wrote the draft. All authors read and approved the final draft.

## Conflict of interest

The authors declare no conflict of interest exists.

## Notes

### Competing Interest Statement

The authors have declared no competing interest.

