## Supplementary material for "P176S polymorphism in Maspin rewires electrostatic interaction that alters Maspin functionality": Suppplementary material

**Prof. Kwang Pyo Kim**

Department of Applied Chemistry, College of Natural Sciences, Kyung Hee University, Yong-in 446-701, Republic of Korea.

### Materials and method

#### Gastric cancer dataset

The proteomic dataset has been downloaded from Clinical Proteomic Tumor Analysis Consortium (CPTAC) data portal (https://cptac-data-portal.georgetown.edu), the sample collection and preparation have been reported previously (Mun *et al.*, 2019).

The mass spectrometry data have been processed using OpenMSv2.5 (Röst *et al.*, 2016) in Konstanz Information Miner (KNIME)v4.2.2 using database matching approach. For the database creation, the human proteome has been downloaded from UniProt with canonical and isoform sequences to be used as the reference data for database matching method. The peptide has been identified using MS-GF+ (Kim and Pevzner, 2014) and OMSSA (Open Mass Spectrometry Search Algorithm) (Geer *et al.*, 2004) search engines and the consensus peptide was retained with and false discovery rate of 1%. The peptide-to-protein identification has been performed using Percolator algorithm (Spivak *et al.*, 2009). Proteins with at least two peptides with >1% threshold has been included for further analysis.

For the peptide identification, precursor mass tolerance was set to 10.0 ppm with a charge range of 1-5, and fragment mass tolerance was 0.3. The cysteine modification (carbamidomethylation), and iTRAQ(isobaric tags for relative and absolute quantitation)-4plex modification of lysine and N-terminal of peptides were used as fixed modifications, while, methionine oxidation and tyrosine-iTRAQ4 modification were the variable modifications. The semi-Trypsin/P was used as the computational digestion protocol. The parameters were largely similar for both search engine, however, for MG-SF+, instrument was set to Q-Exactive, and iTRAQ protocol was used with two maximum modifications. The protein quantification has been performed using IsobaricAnalyzer (Röst *et al.*, 2016) module in OpenMSv2.5 with default isotope correction matrix and normalization.

**Fraction of native contacts**

The fraction of native contacts (Q) has been calculate using MDAnalysis (Michaud-Agrawal *et al.*, 2011) with contacts between heavy atoms with a distance cutoff of 4.5Å and hard cut method was used (Best, Hummer and Eaton, 2013). The residues, θi and θj said to be in contact if the distance between any heavy atoms is less than the cutoff distance and residues are | θi - θj | > 3, then Q is defined as:

$$Q(X) = \frac{1}{N} \sum_{(i,j)} \left( \frac{1}{1 + \exp[\beta(rij(X) - \lambda \mathring{r}ij)} \right)$$

Where; N = total number of paired contacts between atoms “i” and “j”, $r^{0}_{ij}$ is the distance between i and j in the native state, $r_{ij}(X)$ is the distance between i and j in configuration X,

β is a smoothing parameter taken to be 5 $Å^{-1}$ and the factor λ accounts for fluctuations when the contact is formed, taken to be 1.8 for the all-atom model.

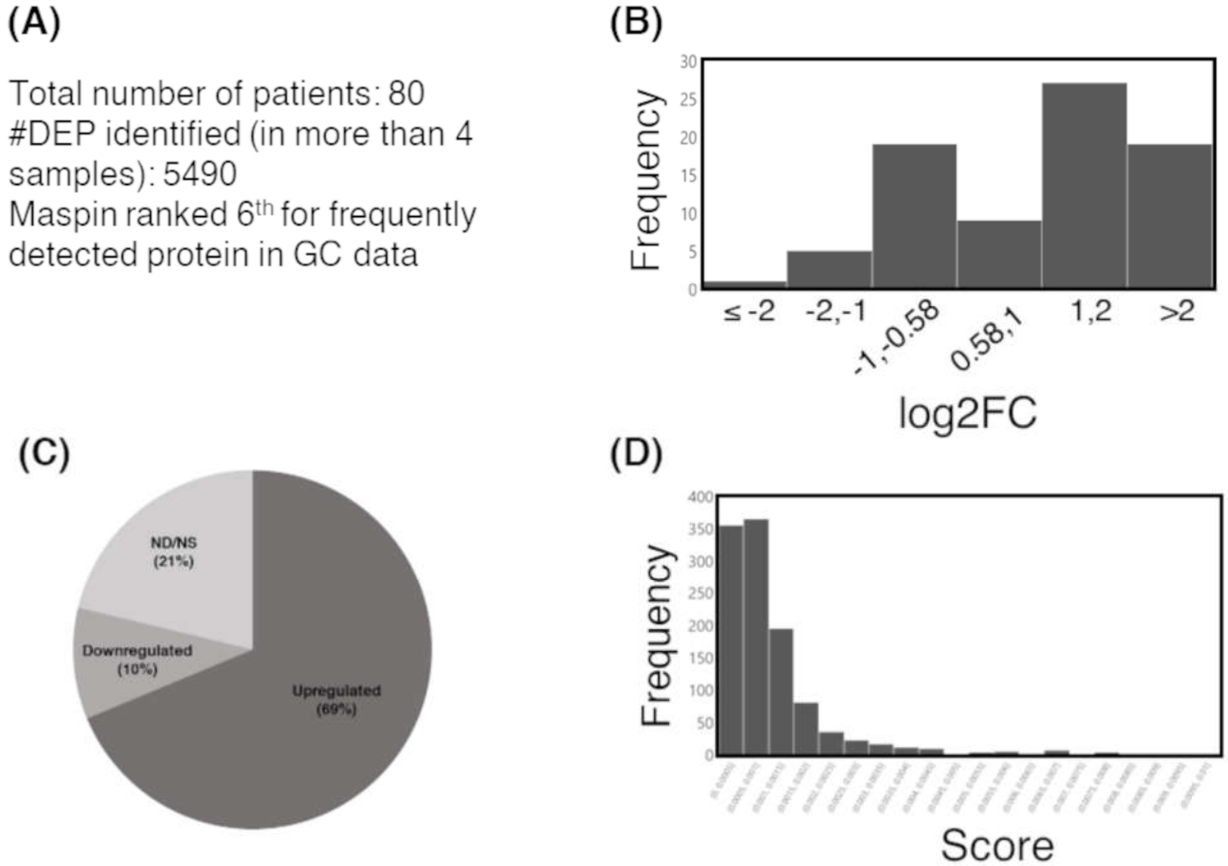


**Supplementary Fig 1. Proteomic data and maspin distribution**. (A) The sample statistics of gastric cancer (GC) data. (B) Distribution of differential expression of maspin in GC dataset. log2Fold change (FC) ≥ ±0.58. (C) The up(down)regulation of maspin in GC dataset. It has been overexpressed in most of the patients. (D) Distribution of protein identification score from OpenMS. DEP, differentially expressed proteins; log2FC, log2 fold change; ND/NS, not detected/not significant.

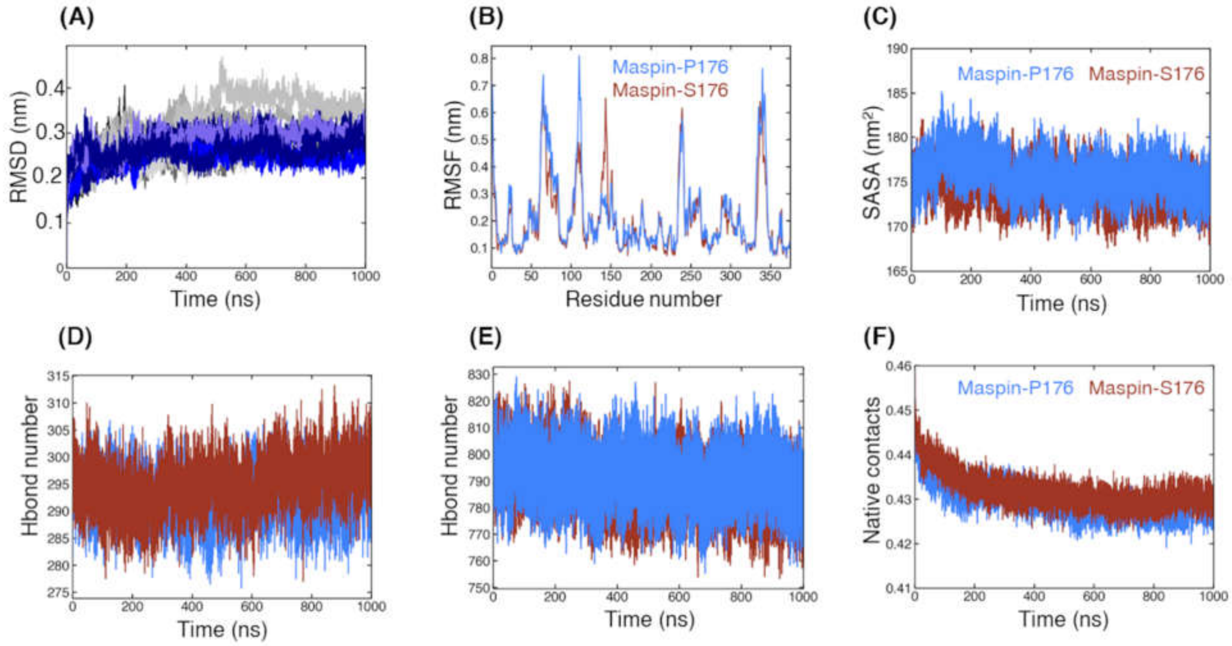


**Supplementary Figure 2: Structural parameters of molecular dynamic simulation of maspins.** (A) The root-mean square deviation (RMSD) values from different simulations. The values in light gray color are for maspin-P176, and different shades of blue refer to maspin-S176. (B) Per-residue fluctuations for the last 500ns of each trajectory. (C) the change in fluctuation over time. (D) Hydrogen bond analysis with 3.5 Å. (E) Number of hydrogen bonds between protein and water (Student T-test p-value=0.626). (F) Fraction of native contacts between the heavy atoms of proteins, with cutoff of 4.5 Å and four residues apart (Student T-test p-value << 0.001).

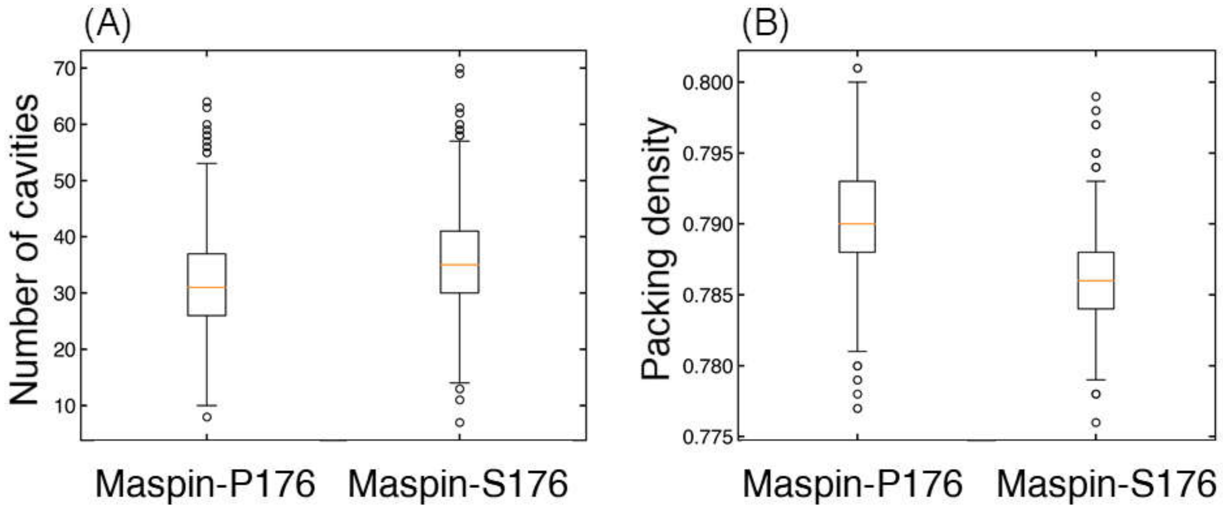


**Supplementary Figure 3: Compactness of maspins.** (A) the number of cavities (B) and the packing density as measured by Voronoia v1.0 with default parameters. For the calculation, 5000 frames from last 500 ns of each trajectory have been used.

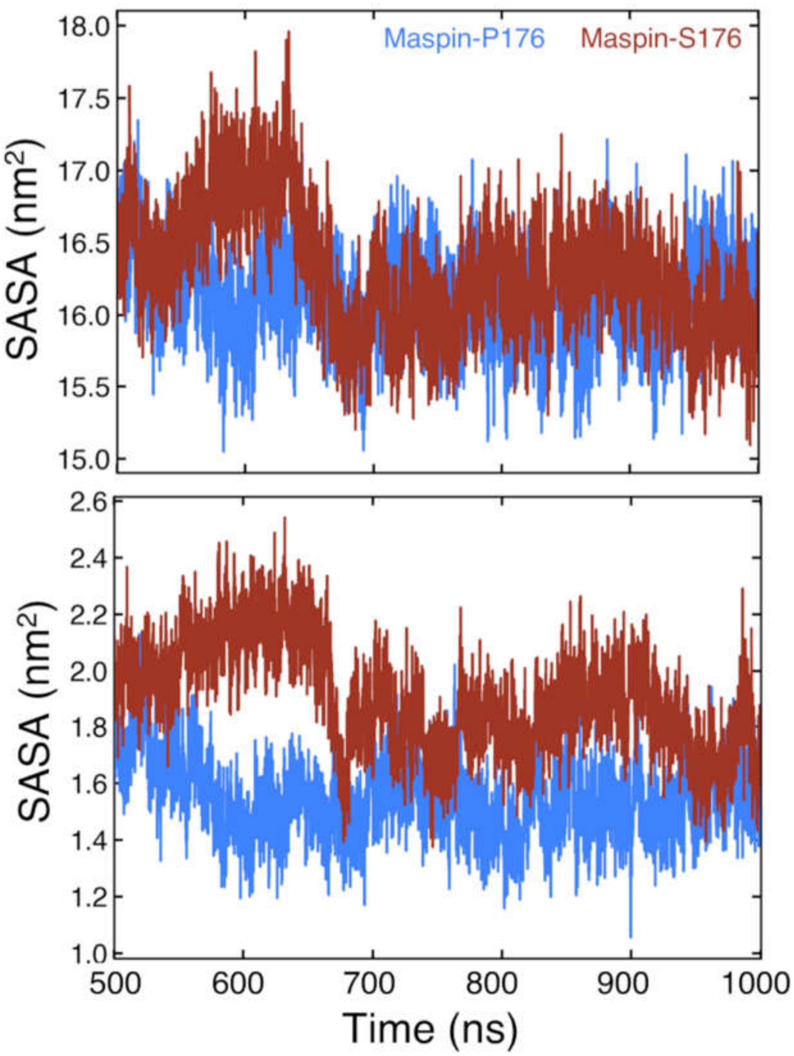


**Supplementary Figure 4.** Solvent exposure of nuclear localization signal (NLS) (region 87-114). (A) The residues from 87-114 have been reported to play role in nuclear localization, thus, solvent accessible surface area (SASA) has been calculated for this region. (B) SASA for 87-95 (s2A in β-sheet A) has been calculated separately.

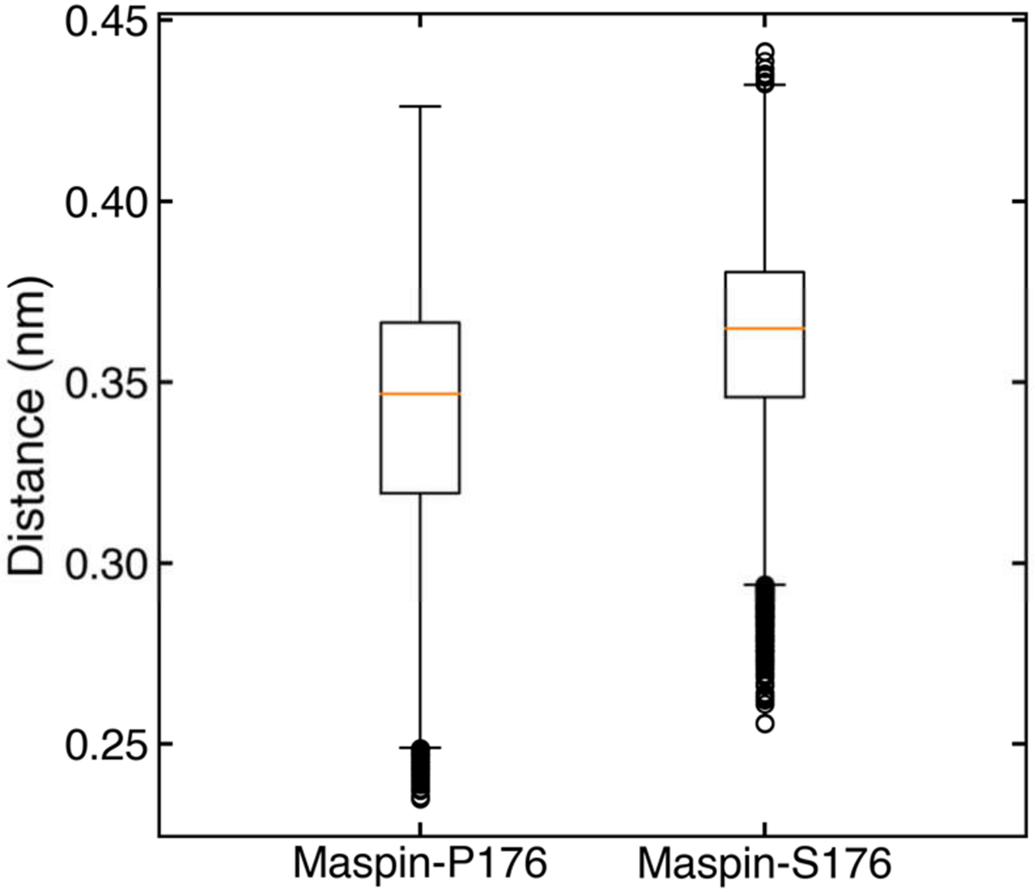


**Supplementary Figure 5: Distance between acidic and basic residues.** The center of mass distance has been calculated for acidic (Asp, Glu) and basic (Lys, Arg, His) residues for the last 500 ns of each trajectory. The box encompasses the first quartile and third quartile, with orange line showing median values of each data. The whiskers are 1.5× the inter-quartile range, and the remaining datapoints are the outliers. Maspin-S176 showed significantly higher distance (Student T-test, p-value << 0.001).

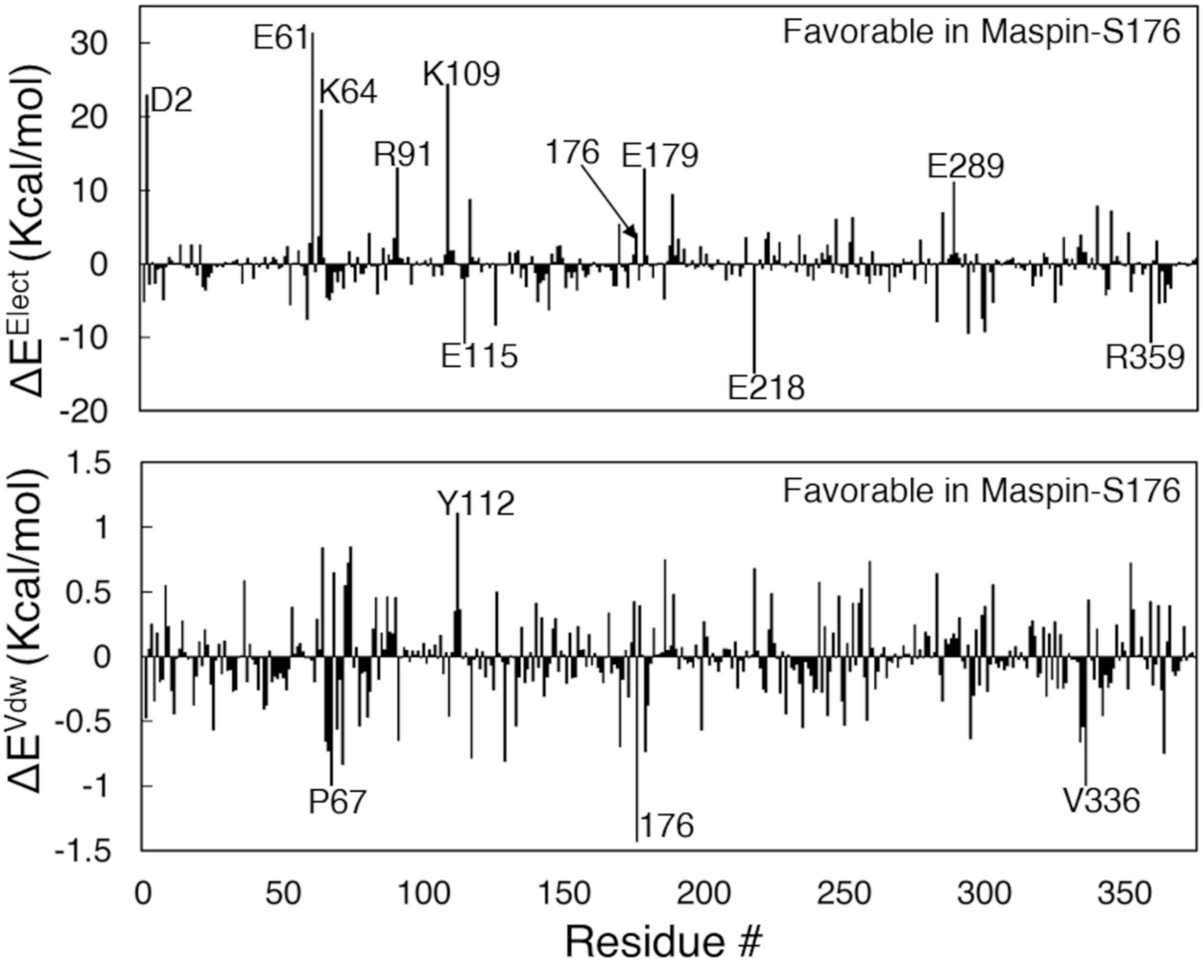


**Supplementary Figure 6. Per-residue energy perturbation between maspin and solvent (water).** The average energy change has been calculated from 5000 snapshots of each trajectory and solvent from two hydration shells have included (2200 water molecules). In this figure, $\Delta E^{Elect} > 10$ and $\Delta E^{Vdw} > 1$ are labelled ($\Delta E^{Elect} = E^{Maspin\text{-}P176}{}_{i,sol} - E^{Maspin\text{-}S176}{}_{i,sol}$).

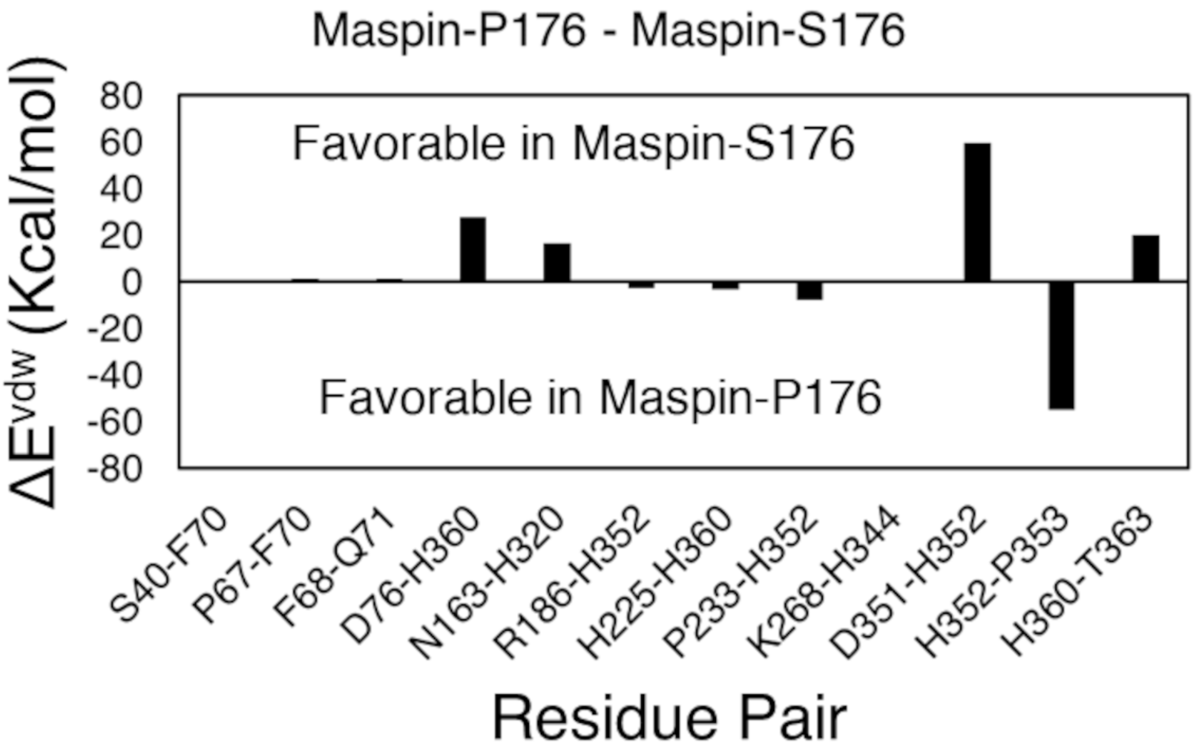


**Supplementary Figure 7. Pair residue $\Delta E^{Vdw}$ interaction energy** with only residue pair with >±1 Kcal/mol are shown.

**Supplementary Table 1:** Differential contacts (in percentage) between maspin-P176 and maspin-S176. Contacts among all residues, for heavy atoms only 4.5 Å cutoff and 4 residues apart (only difference ≥ ±10 are given). The contacts were divided by the highest number of contacts possible in each protein and converted to percentage difference.

| Res1 | Res2 | contacts (Maspin-P176 - Maspin-S176) |
|---|---|---|
| 5 | 76 | -10.36 |
| 8 | 32 | -15.77 |
| 33 | 8 | -15.69 |
| 37 | 70 | -12.49 |
| 40 | 70 | -27.3 |
| 52 | 61 | 15.96 |
| 55 | 61 | 11.42 |
| 60 | 4 | -13.13 |
| 68 | 1 | 10.54 |
| 70 | 60 | 23.61 |
| 84 | 225 | 29.94 |
| 88 | 81 | 10.37 |
| 90 | 115 | -15.76 |
| 90 | 37 | 10.23 |
| 104 | 112 | -11.24 |
| 109 | 104 | -17.86 |
| 129 | 144 | -14.81 |
| 132 | 93 | 10.07 |
| 143 | 166 | 16.55 |
| 144 | 166 | 11.71 |
| 166 | 140 | 10.88 |
| 167 | 84 | -18.27 |
| 168 | 84 | -14.21 |
| 170 | 84 | -23.13 |
| 171 | 219 | 10.52 |

| | | |
|---|---|---|
| 173 | 333 | -12.92 |
| 224 | 335 | -10.15 |
| 229 | 218 | 28.94 |
| 241 | 353 | 17.11 |
| 252 | 10 | 10.88 |
| 259 | 364 | -11.23 |
| 259 | 359 | -11.08 |
| 268 | 344 | -24.09 |
| 268 | 345 | -10.11 |
| 268 | 339 | 11.43 |
| 294 | 289 | 19.81 |
| 297 | 316 | 11.05 |
| 332 | 173 | -15.44 |
| 333 | 172 | -19.75 |
| 340 | 268 | 10.98 |
| 352 | 239 | -13.99 |
| 359 | 218 | -16.76 |
| 359 | 229 | 14.08 |
| 365 | 5 | -17.77 |
| 365 | 73 | 10.23 |

**Supplementary Table 2:** Numerical values of pair residue electrostatic energy between polar residues.

| Residue pair | $\Delta(E_{ij}^{Maspin\text{-}P176} - E_{ij}^{Maspin\text{-}S176)}$ |
|---|---|
| D2-Q5 | -5.2075 |
| D14-H59 | 5.729468 |
| E53-K294 | 6.010128 |
| D76-K362 | 11.91533 |
| K90-E115 | 16.25023 |
| R91-E117 | -5.22796 |
| K109-R110 | 5.568805 |
| K114-E115 | -4.32219 |
| E126-K129 | -7.17416 |
| K137-D141 | 6.7589 |
| K158-E299 | 7.791469 |
| K173-E335 | -4.77479 |
| 176-E179 | 4.698996 |
| E179-K181 | -9.32982 |
| R186-D235 | -5.96681 |
| K189-D238 | -5.59935 |
| K189-E237 | 4.533781 |
| K189-E239 | -4.9788 |
| E201-K345 | 8.414704 |
| K215-K234 | -6.16774 |
| K215-E347 | 4.281948 |
| E218-K224 | -5.88562 |
| E218-S227 | -4.02047 |
| D235-E237 | 5.733291 |
| D238-K245 | -5.84955 |
| E247-K371 | -8.13321 |
| R359-R364 | -4.48217 |

| | |
|---|---|
| K268-E335 | 4.94916 |
| K268-K345 | -4.20518 |
| K270-E347 | -4.40656 |
| E289-K294 | -11.6431 |
| R340-K345 | 5.244918 |

**Supplementary Table 3:** The numerical values of differential allosteric communication intensity (ΔACI) analysis. The values for $\Delta ACI^{(Maspin\text{-}P176 - Maspin\text{-}S176)} > 600$ are given in this table. There were few residues that showed higher coupling intensity in maspin-S176 (indicated by negative values), thus, all residues with negative values are given.

| Residue# | $ACI^{Maspin\text{-}P176}$ | $ACI^{Maspin\text{-}S176}$ | $\Delta ACI^{(Maspin\text{-}P176 - Maspin\text{-}S176)}$ |
|---|---|---|---|
| 16 | 922.9536 | 195.7338 | 727.2199 |
| 24 | 94.34799 | 96.01929 | -1.6713 |
| 29 | 833.3432 | 186.2651 | 647.0782 |
| 65 | 5.482302 | 6.416012 | -0.93371 |
| 67 | 19.88872 | 37.9513 | -18.0626 |
| 109 | 66.25503 | 90.60609 | -24.3511 |
| 121 | 855.8863 | 192.9662 | 662.9201 |
| 167 | 946.3293 | 207.6838 | 738.6455 |
| 170 | 861.2117 | 226.7249 | 634.4868 |
| 171 | 1085.503 | 228.9657 | 856.537 |
| 174 | 1005.925 | 234.1566 | 771.7687 |
| 175 | 1219.444 | 265.0495 | 954.3942 |
| 185 | 1427.148 | 291.1094 | 1136.038 |
| 186 | 1300.678 | 264.9365 | 1035.741 |
| 187 | 1043.988 | 271.9393 | 772.0483 |
| 195 | 988.3001 | 268.6475 | 719.6526 |
| 197 | 1385.796 | 292.7063 | 1093.089 |
| 198 | 1272.716 | 272.189 | 1000.527 |
| 199 | 860.7794 | 239.3888 | 621.3906 |
| 200 | 988.8613 | 248.0089 | 740.8524 |
| 203 | 997.7607 | 203.8742 | 793.8864 |
| 204 | 1169.534 | 260.4973 | 909.0372 |
| 206 | 1191.134 | 255.602 | 935.5321 |
| 209 | 999.7326 | 222.9148 | 776.8178 |
| 212 | 975.3533 | 230.5902 | 744.7631 |
| 214 | 892.8638 | 223.4046 | 669.4591 |
| 216 | 1076.656 | 225.2528 | 851.4031 |
| 217 | 1079.19 | 228.9037 | 850.2865 |

| | | | |
|---|---|---|---|
| 219 | 983.3601 | 222.5592 | 760.8009 |
| 221 | 949.861 | 208.869 | 740.992 |
| 226 | 790.1724 | 175.6681 | 614.5043 |
| 228 | 803.9305 | 194.4639 | 609.4666 |
| 229 | 847.1443 | 187.8933 | 659.251 |
| 230 | 843.8768 | 203.4362 | 640.4406 |
| 231 | 951.7496 | 216.0344 | 735.7153 |
| 232 | 1099.612 | 237.8867 | 861.725 |
| 236 | 844.2617 | 197.351 | 646.9107 |
| 241 | 865.1854 | 216.1294 | 649.0561 |
| 243 | 1078.446 | 225.2955 | 853.1501 |
| 246 | 943.2975 | 220.4906 | 722.8069 |
| 258 | 945.5258 | 200.4601 | 745.0657 |
| 264 | 1032.219 | 235.7598 | 796.459 |
| 269 | 1029.518 | 273.454 | 756.0637 |
| 271 | 1226.122 | 267.2919 | 958.8298 |
| 272 | 959.8644 | 241.5402 | 718.3242 |
| 273 | 1219.87 | 270.3453 | 949.5246 |
| 274 | 1133.476 | 270.8252 | 862.6508 |
| 322 | 844.8757 | 194.7044 | 650.1713 |
| 326 | 922.6967 | 230.088 | 692.6086 |
| 327 | 895.6658 | 226.5035 | 669.1623 |
| 329 | 854.9385 | 221.4293 | 633.5092 |
| 330 | 1112.287 | 244.4097 | 867.8773 |
| 348 | 923.2454 | 238.1186 | 685.1268 |
| 350 | 1108.992 | 255.6129 | 853.3794 |
| 352 | 999.7834 | 262.8928 | 736.8905 |
| 353 | 1159.962 | 252.14 | 907.8224 |
| 354 | 1039.697 | 227.2903 | 812.4069 |
| 355 | 1000.645 | 215.9706 | 784.6745 |
| 356 | 939.6398 | 191.2918 | 748.348 |
| 369 | 880.5902 | 181.8333 | 698.7569 |

| | | | |
|---|---|---|---|
| 370 | 877.0177 | 186.3478 | 690.6699 |
| 371 | 872.4034 | 220.2312 | 652.1721 |
| 372 | 1142.647 | 243.1637 | 899.4829 |
| 373 | 1128.229 | 245.3692 | 882.8596 |
